# Visualizing tomato spotted wilt virus protein localization: Cross-kingdom comparisons of protein-protein interactions

**DOI:** 10.1101/2023.11.27.568851

**Authors:** K.M. Martin, Y. Chen, M.A. Mayfield, M. Montero-Astúa, A.E. Whitfield

## Abstract

Tomato spotted wilt virus (TSWV) is an orthotospovirus that infects both plant and insect cells. Understanding the protein localization and interactions in these cells is crucial for unraveling the infection cycle and host-virus interactions. In this study, we investigated the localization of TSWV proteins in cells of plants and insects. Furthermore, we identified the protein-protein interactions among TSWV proteins using bimolecular fluorescence complementation (BiFC) and yeast two-hybrid (MbY2H) assays. Our results revealed distinct localization patterns for TSWV proteins in plant and insect cells. The nucleocapsid protein (N), essential for genome encapsidation, was found in the cytoplasm of both cell types. The non-structural movement protein (NSm) localized to the cytoplasm in insect cells, different from the localization in plant cells’ plasmodesmata. The non-structural silencing protein (NSs) exhibited peripheral localization in plant cells and cytoplasmic localization in insect cells. Additionally, glycoproteins G_N_ and G_C_ showed cytoplasmic localization in both cell types. Moreover, protein-protein interaction analyses revealed self-interactions of NSm, N, G_N_, G_C_, and NSs. These interactions are crucial for viral genome encapsidation, virion assembly, and RNA silencing suppression. We also identified interactions between different TSWV proteins, indicating their roles and host interactions. Comparisons with other orthotospovirus interaction maps highlighted the uniqueness of TSWV protein-protein interaction networks. Despite sharing genome organization and putative gene annotations, each orthotospovirus exhibited distinct interaction maps. Overall, this research expands our knowledge of TSWV infection and elaborates on the intricate relationships between viral proteins, cellular dynamics, and host responses. These findings lay the groundwork for future studies on the molecular mechanisms of TSWV infection and may facilitate the development of effective control strategies.

## Introduction

Tomato spotted wilt virus (TSWV, *Orthotospovirus tomatomaculae)* has a global distribution, this virus infects over 1000 plant species in over 85 families and is considered one of the top ten most devastating plant viruses worldwide (Parrella, 2003; Scholthof et al., 2011). Infected plants include peanuts, tobacco, tomato, bean, spinach, and lettuce among many others causing food security issues in both developed and developing countries (Oliver and Whitfield, 2016). Symptoms of TSWV include necrosis, leaf deformations such as cupping and curling, and concentric chlorosis on fruits depending on the plant species. Crop losses to this virus are estimated to cause over $1.4 billion dollars in damage in a ten-year period in the US alone (Culbreath et al., 2003; Pappu et al., 2009; Riley et al., 2011; Mandal, 2012)It is vectored in a persistent, circulative manner by nine species of thrips with the most efficient being the western flower thrips, *Frankliniella occidentalis.* (Whitfield et al., 2005; Whitfield et al., 2015; Rotenberg and Whitfield, 2018) Currently, the control of orthotospoviruses relies heavily on the management of vector thrips, however, populations of *F. occidentalis* have also developed resistance to many insecticides (Reitz, 2009). Development of alternate, non-traditional means of controlling either the thrips acquisition of viruses or the overall population of the insect vectors is a possible long-term strategy for managing this pathosystem (Pappu et al., 2009; Mandal, 2012).

Characterizing the protein-protein interactions amongst viral proteins is a first step in the identification of factors involved in the virus life cycle. TSWV has a segmented ambisense RNA genome with a small (S), medium (M) and large (L) component. The L component encodes an RNA-dependent RNA-polymerase necessary for viral replication. The M component encodes a polyprotein of two glycoproteins, G_N_ and G_C_, which are cleaved during processing and are part of virion recognition on the insect cell membrane, and a non-structural protein called NSm, thought to be involved in viral movement from cell to cell in plants (Kormelink et al., 1994; Whitfield et al., 2004; Feng et al., 2016). The S component encodes two proteins, a nucleocapsid protein (N) and a non-structural protein called NSs that is involved in silencing of the host defenses (Adkins, 2000). In previous studies, an association between N and NSm has been identified (Soellick et al., 2000), as well as associations with N with itself and Nsm with itself in protoplasts and yeast (Storms et al., 1995; Uhrig et al., 1999). G_N_ and G_C_ have also been shown to interact *in vivo* in both BHK21 cells and plant cells (Snippe et al., 2005; Ribeiro et al., 2009). However, a comprehensive protein-protein interaction map has not been created for TSWV in the same systems under similar conditions. To better understand the virus infection cycle in insect and plant hosts, the associations of N, NSs and NSm to the glycoproteins or to NSs need to be determined. Identification of conserved interactions between viral proteins will inform future work on targets for disruption of the viral replication and movement as another means of control of these plant pathogens. It will also further guide studies on the possible involvement of critical host factors necessary for viral success in the host.

As TSWV infects both a primary host and arthropod vectors, both hosts present opportunities for understanding the virus replication cycle and for interdiction. The goals of this project were to i) create a protein-protein interaction map of TSWV proteins in plants and insect cells and ii) compare the TSWV protein localization patterns in the absence and presence of viral infection *in planta.* This study is the first comparative analysis of protein localization for an orthotospovirus in both animal and plant cells. This work provides a platform for the identification of molecules that disrupt the protein interactions in both the vector and plant hosts.

## Materials and Methods

### Plant growth and maintenance

Wildtype and transgenic *N. benthamiana* expressing cyan fluorescent protein fused to histone 2B (CFP:H2B) were grown in either a growth chamber set at 25°C for 14 hours of daylight at 300 µmol/m2/sec light intensity and a 10-hour dark period, or in a greenhouse under ambient conditions. Plants were transplanted from seedling pots to individual containers at two weeks of age. After transplanting, plants were allowed to grow for approximately one to two weeks further until the leaves reached the size of quarters. This size corresponded to the best age for either the infiltration of plants for microscopy or the inoculation of plants with virus described below.

### Inoculation of plants with TSWV

*Emilia sonchifolia* plants were inoculated with TSWV following the protocol described (Ullman et al, 1992, Ullman et al, 1993). Once symptoms developed, they were utilized as the inoculative source to mechanically inoculate *N. benthamiana* with 300 grit Silicon Carbide and 10mM Sodium Sulfite. Once symptoms were first visible at approximately 7-9 days in summer and 9-11 days in fall and winter months post-inoculation, plants were infiltrated with TSWV constructs for protein expression in the manner described in Badillo-Vargas et al., 2019. The point in which the leaves were infiltrated was deemed an “early infection point” as the leaves began to show a mild mosaic and a slight leaf curl indicative of infection compared to a more advanced state with severe leaf curling and more dramatic mosaic patterns. At the later time point, infiltration becomes difficult due to leaf architecture changes including leaf thinning and curling which causes difficulties in infiltration.

### Identification of a cellular marker to indicate infection with TSWV

A cellular marker for infection is important in a systemic infection as the leaves are infiltrated just prior to a full infection. In this case, some cells may remain healthy compared to others that are infected. *Nicotiana benthamiana* plants with transgenic markers for red histone-2B (RFP-H2B) and red endoplasmic reticulum markers (RFP-ER) were inoculated with TSWV with similar methods as described above. After symptoms were prevalent in systemic non-infiltrated leaves at between 7-11 days depending on season, leaf samples were cut and utilized for confocal microscopy to observe cellular changes due to the infection. Additionally, *N. benthamiana* plants were infected with TSWV and infiltrated with unfused red fluorescent protein when leaves were identified as an early infection point of the leaves as described earlier. Leaves that were rub inoculated with virus are not imaged due to leaf damage during the inoculation.

### Cloning of TSWV genes for expression in plants and insects

Cloning each of the TSWV open reading frames for expression represents the first step to test both localization and potential interactions. TSWV-N was amplified from pBS-N2C4.5 (Kim et al., 1994) using primers, F: 5’ CACCATGTCTAAGGTTAAGCTCACTAAGG 3’ and R: 5’ AGCAAGTTCTGTGAGTTTTGC CTG 3’. TSWV-NSs was amplified using primers, F: 5’ CACCATGTC TTCAAGTGTTTATGAGTCG 3’. TSWV-NSm was amplified in two forms, one with a stop codon using primers, F: 5’ CACCATGTTGACTCTTTTCGGTAATAAGG 3’; R: 5’ TTATATTTCATCAAAAGACAACTGAGCAACACTG 3’. A second form of NSm without a stop codon was amplified with the same forward primer and the reverse primer: 5’ TATTTCATCAAAAGATAACTGAGCAAC 3’. G_N_ was also amplified from pGF7 (Adkins et al., 1996) in two forms, one full-length protein, Forward, 5’ CACCATGAGAATTCTAAAACTACTAGAACTAGTCG 3’ and Reverse, 5’ CATAGACATGGGCATTTGAGACAAAATGATC 3’ and a soluble form without a transmembrane domain with the same forward primer and the reverse primer: 5’ AATGCTTTTTGAATATTTGATTATGCAATCTCTAAC 3’. The G_c_ was also amplified from pGF7 (Adkins et al., 1996) in two forms, one full length and one soluble form without the transmembrane domain. The full-length form primers were forward, 5’ CACCATGTTGATCATTTTGTCTCAAATGCCC 3’ and reverse, 5’ AAGCCACCTATGGATTTCTCTCACCTT GTC 3’. The soluble form was amplified using the same forward primer as the full-length version and the reverse primer: 5’ ATAGCTTGCAATGAAATTGAATGGGC 3’. Once the open reading frames (ORF) for the TSWV genes stated above were amplified, they were cloned into pENTR-d TOPO (Thermo Fisher Scientific) following manufacturer’s instructions. Once cloned, the plasmids were sent for sequencing to confirm the identity of each ORF.

### Transfer of TSWV ORFs into expression vectors for plants

To express each of the TSWV ORFs in plants and determine localization, entry clones for each of the TSWV ORFs generated previously (Badillo-Vargas et al., 2019) were moved into plant expression vectors. This was done using LR clonase II (Invitrogen) into the pSITE-EGFP-C1 vector or pSITE-EGFP-N1 (Chakrabarty et al., 2007) following the manufacturer’s protocol provided. To express each of the TSWV ORFs to determine protein-protein interactions with bimolecular fluorescence complementation assays (BiFC) between the viral proteins, TSWV entry clones were moved into pSITE-NEC, pSITE-CEC, pSITE-NEN or pSITE-CEN (Martin et al., 2009) using LR clonase II (Invitrogen) using the manufacturer’s protocol. All expression vectors were confirmed with sequencing.

### Expression of TSWV proteins in plants

TSWV ORFs N, NSs, NSm, G_N_ and G_C_ were cloned in frame to green fluorescent protein (GFP) as described above and were transformed into *Agrobacterium tumefaciens* strain LBA 4404 and used for transient infiltrations of *N. benthamiana.* To determine their cellular localization, a cellular marker was used, the nuclear marker (Histone H2B) fused to red fluorescent protein (Martin et al., 2009). These were also co-infiltrated with unfused RFP into wildtype *N. benthamiana* either healthy or TSWV symptomatic plants (described previously) of the same age. TSWV proteins fused to GFP were localized at 2 days post infiltration. TSWV-GFP constructs were also infiltrated into healthy cells and visualized at both 2 and 3-day time points after infiltration in healthy plants. To test protein-protein interactions, expression clones for TSWV ORFs were generated in context for YFP halves (described previously) were transformed into *Agrobacterium tumefaciens* stain LBA 4404. These constructs were pooled in all pairwise combinations and infiltrated into *N. benthamiana* expressing the nuclear marker cyan fused to histone 2B (Martin et al., 2009) following the procedures described in Badillo-Vargas et al., 2019. BiFC constructs were imaged at both two days and four days after infiltration. Each pairwise combination was infiltrated no fewer than three times, capturing at least three independent cells during each microscopy event. Each interaction was also compared to a protein-non-binding control infiltration to ensure that the fluorescence of the interacting partners was above the background seen when the YFP halves may come together by chance. This also helped to eliminate interactions that were not specific but demonstrated a protein’s ability to bind to any other protein. GST was selected as a non-binding control in plants as it was used previously for virus-virus protein interaction mapping for both tospoviruses and rhabdoviruses (Martin et al., 2009; Min et al., 2010; Dietzgen et al., 2012; Martin et al., 2012; Widana Gamage and Dietzgen, 2017; Martin and Whitfield, 2018). For combinations where the comparison to the non-binding controls resulted in confounding results, no less than six separate infiltrations were performed with at least three cells captured each time to ensure that the interaction was above background.

### Expression of TSWV proteins in Sf9 insect cells

TSWV-ORFs fused to sequences encoding fluorescent proteins were constructed for localization in insect cells. pENTR/D-TOPO (Thermo Fisher Scientific) entry clones containing individual TSWV proteins were recombined with a pIB/V5-His destination vector (Thermo Fisher Scientific) containing either GFP or mRFP in both amino and carboxy terminal fusions when possible. The resulting constructs were pIWG or pIWR vectors (gateway cassette before the fluorescent protein) and pIGW or pIRW vectors (gateway cassette after the fluorescent protein) (Teh et al., 2022). To determine if TSWV proteins share a similar localization pattern in insect cells to plant cells, lepidopteran ‘SF9’ cells were transfected at approximately 85-90% confluency with plasmids containing an individual TSWV protein fused to GFP in a 35 mm^2^ dish using Cellfectin II (Gibco) following manufacturer’s recommendations. Transfected cells were incubated at 28°C for 72 hours. Each transfection also contained a positive control of pHSP-70-GFP to determine if the transfection itself was successful (Martin and Whitfield, 2018), as well as a negative control which contained no plasmid DNA to eliminate any internal cell fluorescence. Each construct was localized between 5-20 times for microscopy photos, obtaining 3 photosets from each localization during replication.

### Microscopy of plant and insect cells

Sections of plant leaf tissue between two days to four days after infiltration were mounted in water on a microscope slide to prepare for microscopy. All images were acquired on a Ziess LSM 780 laser scanning confocal microscope using the C-Apochromat 40x/1.2 W Korr M27 objective. Image acquisition was conducted on Zen 2 black edition v. 10.0.0 at 1024 x 1024 pixels with a scan rate of 1.58μs per pixel, 16 bit depth, and with pixel average of 4. Zen 2 blue edition lite 2010 v. 2.0.0.0 was used for image conversion to jpeg format. Figures were assembled using Microsoft Powerpoint 2008 version 12.8.6. Microscopy of insect cells was performed at 72 hours post-transfection using an Eclipse Ts2R (Nikon) epifluorescent microscope using an S Plan Fluor ELWD 40x/.060 objective with a CFI 10x/22 eyepiece. Image acquisition was conducted using Nikon Elements software v. 5.21.02 (Build 1487). Each of the nine (or more) pictures were compared to each other during figure assembly to ensure the best representative image was used to capture the described results. Figures were assembled using Microsoft Powerpoint v. 2305 (Build 16501.20210 Click-to-Run).

### Validation of interactions between TSWV proteins using the split-ubiquitin membrane-based yeast two-hybrid system

The split-ubiquitin membrane-based yeast two-hybrid (MbY2H) system was used for identifying protein interactions with the viral glycoproteins, G_N_ and G_C_, due to their transmembrane domains. Two-step transformation was performed for MbY2H, the first step was to transform each recombinant pBT3-SUC plasmid containing a TSWV gene into NYM51 yeast (*Saccharomyces cerevisiae*) competent cells. Competent cell preparation and transformation were performed based on manufacturer’s protocol and previously described methods (Dualsystems Biotech, Schlieren, Switzerland, Badillo-Vargas et al., 2019). For each transformation reaction, 50 µg of denatured carrier DNA and 1.5 µg of plasmids were added into 100 µl freshly prepared competent cells. Polyethylene glycol-lithium acetate (PEG/LiAc) and dimethyl sulfoxide (DMSO) were used for transformation. After incubation and transformation, cells were recovered in 1 ml of yeast-peptone-dextrose (YPD) medium for 1.5h. Then, cells were pelleted and resuspended in 100 µl 0.9% NaCl and spread onto SD/–Trp agar media. The colonies expressing individual TSWV-Cub fusion proteins were then used to make competent cells, and the second step was to transform each recombinant pPR3N-TSWV construct (1.5 µg plasmid/transformation reaction) into the yeast expressing TSWV-Cub. Interactions among TSWV proteins were validated by interactions among fusion protein pairs, TSWV-Cub and NubG-TSWV. Interactions between TSWV-Cub and NubI (transformation of pOst-NubI into TSWV-Cub expressing NYM51 cells) and between TSWV-Cub and NubG (transformation of pPR3N empty vector into TSWV-Cub expressing NYM51 cells) were used as positive and negative controls, respectively. Co-transformation of pTSU2-APP and pNubG-Fe65 into NYM51 was used as another positive control. All transformants were cultured on both SD/–Leu/–Trp double-dropout (DDO) (Supplemental Figure 2) and SD/–Ade/–His/–Leu/–Trp quadruple-dropout (QDO) (Figure 4) media at 30°C. The entire experiment was performed three times, each experimental replicate included three transformation replicates of individual interactions. Colonies grown on QDO media that represented the potential positive interactions were randomly selected and confirmed by *β*-galactosidase assay following the manufacturer’s protocol (Thermo Fisher Scientific, (Badillo-Vargas et al., 2019)).

### Controlling auto-activation in the split yeast two-hybrid system

Due to auto-activation of Gc-Cub, 3-amino-1,2,4-triazole (3-AT) was added to the QDO media to optimize the screening stringency of G_C_ using positive and negative controls. 1.5 µg of plasmids pPR3N (negative control) or pOst-NubI (positive control) were transformed into 100 µl competent cells expressing Gc-Cub, the transformants were resuspended in 200 µl 0.9% NaCl and 100 µl was spread onto both DDO and QDO agar media supplemented with different 3-AT concentrations. The addition of 100mM 3-AT suppressed the growth of the negative controls but not positive controls, and was used for validating G_C_-Cub and Nub-TSWV interactions.

## Results

### Localization of TSWV proteins in healthy cells of plants and insects

The localization of the TSWV viral proteins in plant and insect cells was conducted to determine the similarities and differences when the virus replicates in cells from different kingdoms. When expressed in insect cells, NSm was found surrounding the nucleus, as opposed to plasmodesmata in plant cells. N and NSs were found in the cytoplasm of insect cells, while N is found in the cytoplasm for plants and NSs is found in the cellular periphery of plant cells. G_N_ localizes in insect cells in the cytoplasm outside of the nucleus. The localization of a soluble G_C_ (G_C_S, lacking the transmembrane domain and cytoplasmic tail) appears to mirror G_N_, also localizing in the cytoplasm surrounding the nucleus (Figure 1 XVI). Localization of the full length G_C_ was conducted in marker plants and insect cells several times (minimum of 7 times for each cell type) and the protein was not detected with consistency. It was localized in the presence of unfused RFP as a transient marker in plants (described below); however, localization of Gc was challenging and to achieve our minimum standard of three replicated localizations, thus we had to perform more than twenty independent infiltration experiments.

**Figure 1.**
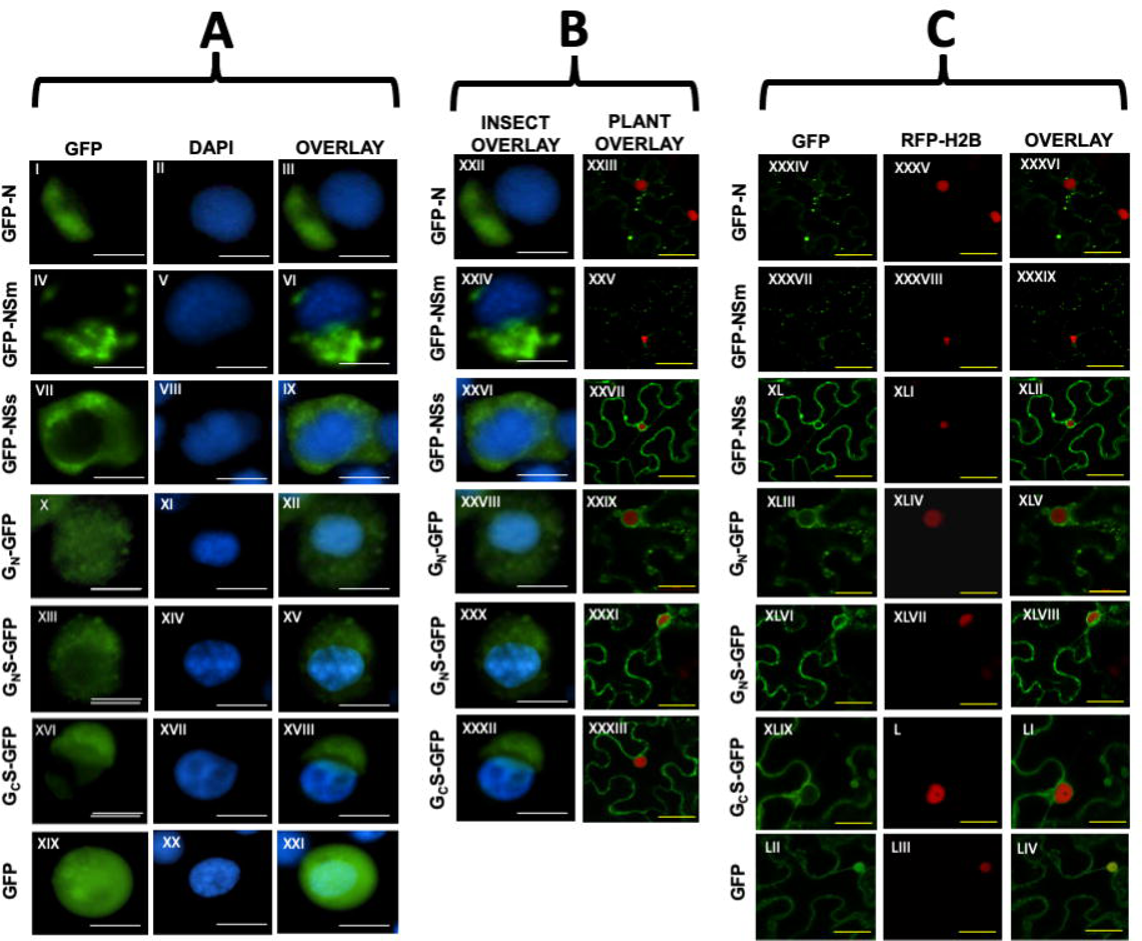
Localization of TSWV proteins in insect and plant cells. A) TSWV proteins expressed in SF9 cells. Column 1, GFP tagged to TSWV protein, Column 2, DAPI stain to indicate nucleus, Column 3, overlay of Column 1 and 2. Rows are as follows: I-III, TSWV-GFP-N, IV-VI, TSWV-GFP-NSm, VII-IX, TSWV-GFP-NSs, X-XII, TSWV-G_N_-GFP, XIII-XV, TSWVG_N_S-GFP, XVI-XVIII, TSWV-G_C_S-GFP, XIX-XXI, Control. B) Comparison of overlay for both SF9 cells and plant cells. Column 1, overlay of insect cells, Column 2, overlay of plant cells. Rows are as follows: XXII-XXIII, TSWV-GFP-N, XXIVXXV, TSWV-GFP-NSm, XXVI-XXVII, TSWV-GFP-NSs, XXVIII-XXIX, TSWV-G_N_-GFP, XXX-XXXI, TSWV-G_N_S-GFP, XXXII-XXXIII, TSWV-GcS-GFP. C) TSWV proteins expressed in plant cells. Column 1, GFP tagged to TSWV protein, Column 2, RFPH2B to indicate the nucleus of the cell, Column 3, overlay of columns 1 and 2. Rows are as follows: XXXIV-XXXVI, TSWV-GFP-N, XXXVII-XXXIX, TSWV-GFP-NSm, XL-XLII, TSWV-GFP-NSs, XLIII-XLV, TSWV-Gn-GFP, XLVI-XLVIII, TSWV-G_N_S-GFP, XLIX-LI, TSWV-G_C_S-GFP, LII-LIV, Control. White scale bars (insects) indicate 25μm while yellow scale bars (plants) indicate 20μm.

### Localization of tomato spotted wilt virus proteins in plants over time

Previous experiments in plants indicated that TSWV protein localization changes over time post-agroinfiltration (Montero-Astúa, 2012). At an early time point, two days after infiltration, N was present on the cellular periphery and in aggregations in the cell. After one additional day, three days post-infiltration, N accumulated to a greater amount in the interior of the cell with little seen on the periphery (Montero-Astúa, 2012). At two days post-infiltration, NSs localized along the cell periphery and NSm was present both at the cell periphery and in aggregations in the cell (Supplemental Figure 1). The protein localization moved to the interior of the cell with both NSs and NSm at three days post-infiltration as well (Supplemental Figure 1). No changes were observed over time with the localization of G_N_ and G_C_ (data not shown). In insect cells, the localization pattern of viral proteins did not change over time, however, the fluorescence intensity increased (data not shown).

### Identification of a cellular marker to indicate infection with tomato spotted wilt virus

It was of interest to determine if localization changes occurred during viral infection compared to the single infiltrations of proteins in healthy cells. Changes in the localization could indicate possible interactions and movements which only occur in the presence of other viral proteins, such as protein complexes and the L protein which we did not express in this study. It was then important to identify a cellular marker that would identify when cells were infected with virus, because we needed to infiltrate into *Nicotiana benthamiana* plants prior to the full infection when virus may have a patchy distribution and no clear symptoms (Goodin et al., 2007). This is due to the inability to infiltrate mature-fully infected plants effectively and see localization of the protein fusions with *Agrobacterium tumefaciens* transient transformations. As a result, some of the infiltrated cells were infected with TSWV and in some cases the viral infection front would not have reached the site of infiltration and the cells were healthy at two days post-infiltration. In plants with TSWV and unfused RFP, the RFP formed areas where voids existed that did not exist in the healthy comparisons. These voids appeared as bubbles and were present on the cell periphery or near the nucleus. The unfused RFP was present throughout the cell providing both nuclear and cytoplasmic references and was selected as an additional marker for this reason. There was no single site where there appeared to be a selective preference for these voids to be present as they could be observed in both the cell periphery in various areas and near the nucleus at random. No changes were observed in the RFP which was found in the nucleus (Figure 2). No cellular changes were observed in the RFP-H2B or RFP-ER plants to indicate infection (data not shown).

**Figure 2.**
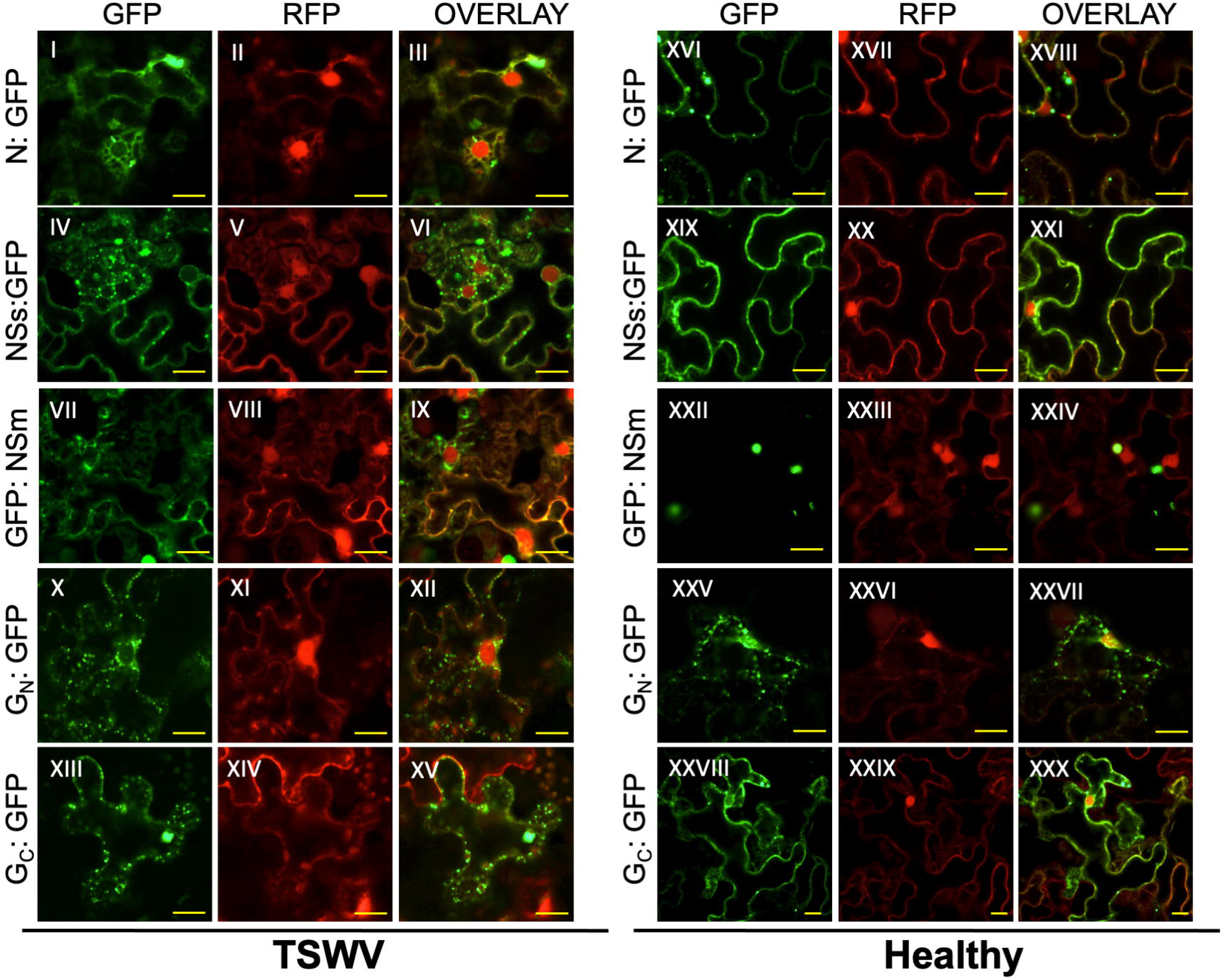
Localization of TSWV proteins with infected and healthy plant cells. Left side) TSWV proteins expressed in agro-infiltrated plant cells. Column 1, GFP tagged to TSWV protein, Column 2, unfused RFP, Column 3, overlay of columns 1 and 2. Rows are as follows: I-III, TSWV-N-GFP, IV-VI, TSWV-NSs-GFP, VII-IX, TSWV-NSm-GFP, X-XII, TSWV-G_N_-GFP, XIII-XV, TSWV-G_C_-GFP. Right side) TSWV proteins expressed in healthy plant cells. Column 1, GFP tagged to TSWV protein, Column 2, unfused RFP; Column 3, overlay of columns 1 and 2. Rows are as follows: XVI-XVIII, TSWV-N-GFP, XIX-XXI, TSWV-NSs-GFP, XXII-XXIV, TSWV-NSm-GFP, XXV-XXVII, TSWV-G_N_-GFP, XXVIII-XXX, TSWV-G_C_-GFP. Scale bar is equal to 20 μm.

### Localization of TSWV protein fusions in both healthy and virus infected cells

As previously indicated, the cellular changes to indicate infection were a bubble type localization pattern of the RFP. This was utilized as a cellular marker to image the TSWV proteins during infection. Because the viral infection front may still be progressing through the infected tissue, some cells appeared healthy even in the visibly infected plants and so only those areas that showed these voids were imaged. TSWV-N GFP localized around the unfused RFP areas and appeared the most soluble during infection compared to when not infected shown on the right side of Figure 2. The N aggregated much more in healthy tissues (Figure 2 XVI) and was more soluble in infected tissues (Supplementary video 1). NSs-GFP is also around the RFP areas, but it was also present in punctate bodies in these areas as well. These tiny NSs bodies were mobile and were not unique to infection but were much more prominent all throughout the cell during infection (Supplementary video 2). GFP-NSm localized to the areas marked by RFP and also formed large aggregations however, in infected tissues, the large aggregations were much less common, compared to many in healthy tissues (Figure 2 XXII, Supplementary video 3).

The viral glycoproteins, G_N_-GFP and G_C_-GFP, also both localized around the red areas that marked TSWV infection and they appeared mostly as small bodies with a similar distribution and shape. This glycoprotein distribution was unique to TSWV-infected cells. G_N_ formed these bodies in both healthy and infected cells. G_C_-GFP, in its full-length form, including the transmembrane domain and cytoplasmic tail had a much more fluid appearance in healthy cells compared to punctate loci seen in infected cells (Figure 2 XXV and XXVIII, Supplementary video 4). The healthy control and virus infected cells were all imaged at two days after infiltration for all tested viral proteins due to the high incidence of the death of the cells in infiltrated virus infected cells. This is thought to be a response to the plant cells potentially overwhelmed by the expression of both transient viral proteins and an active viral infection.

### Bimolecular fluorescence complementation (BiFC) assays

To identify the possible protein-protein interactions between TSWV proteins associated with the changes in localization during infection, BiFC analysis was done. Four positive interactions were identified using BiFC. TSWV N self-interacted and this interaction was localized to the cell periphery. The interaction was more prevalent at 4 dpi, but also seen at 2dpi. The NSm protein also self-interacted and was localized to the cell periphery. The localization pattern was small-punctate bodies where the proteins interacted. This was different from the overall localization of the GFP-NSm fusion (Figure 2 VII) suggesting the interaction is very specific in location as seen in the GFP (Figure 2 LII).

There were two protein-protein interactions which occurred more reproducibly at one of the two timepoints observed in the BiFC experiments. The NSs and N interaction was reproducibly seen at 4 dpi (Figure 3, XXII-XXIV). Although it could be seen irregularly at 2 dpi, the N-NSs interaction was not considered above the level of background fluorescence at this early time point. Consistent with the N self-interaction at 4 dpi, there was a greater amount of N present in internal areas, the NSs and N interaction changes may reflect the increase in N expression or accumulation. The second interaction which was seen regularly at one time point was NSs and GcS which was present only present at the cell periphery, not near nucleus and only seen at 2dpi (Figure 3 X-XII). The NSs-GcS interaction could not be visualized reproducibly above background at 4 dpi (Figure 3, XXV-XXVII).

**Figure 3.**
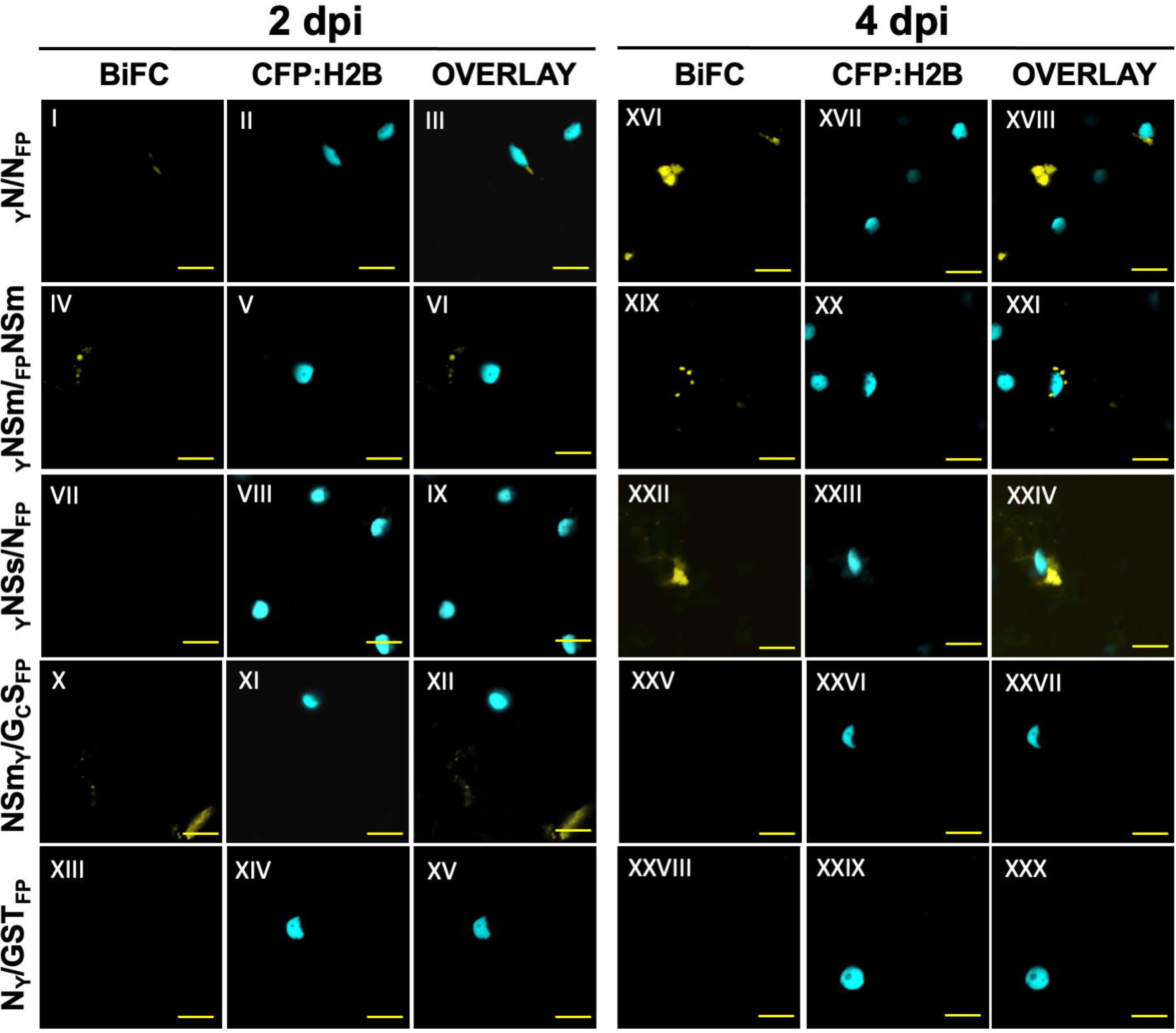
Bimolecular Fluorescence complementation map of tomato spotted wilt virus. From left to right: column 1 is the YFP channel to detect possible BiFC at 2dpi, column 2 is the CFP channel for detection of the nuclear marker histone 2B (CFP:H2B) at 2dpi, column 3 is the overlay of columns 1 and 2, column 4 is the YFP channel to detect possible BiFC at 4dpi, column 5 is the CFP channel for detection of the nuclear marker histone 2B (CFP:H2B) at 4dpi, and column 6 is the overlay of columns 4 and 5. Row 1, I-III + XVI-XVIII, TSWV-N fused to the YFP amino terminal-half with TSWV-N fused carboxy half of YFP. Row 2, IV-VI + XIX-XXI, TSWV-NSm fused to the amino terminal half of YFP with NSm fused to the to the carboxy half of YFP. Row 3, VII-IX + XXII-XXIV is TSWV-NSs fused to the amino terminal half of YFP with TSWV-N fused to the carboxy half of YFP. Row 4, X-XII + XXV-XXVII, TSWV-NSm fused to the amino terminal half of YFP with TSWV-G_C_S fused to the carboxy half of YFP. Row 5, XIII-XV + XXVIII-XXX, TSWV-N fused to the amino terminus of half of YFP with Glutathione S-Transferase fused to the carboxy terminus of YFP. Scale bar is equal to 20 μm

### Interactions of TSWV proteins with each other tested via membrane-based yeast two-hybrid analyses

To further characterize TSWV protein-protein interactions, we used the MbY2H system to test all viral proteins for possible interactions, except the L protein. Due to the different predicted cleavage sites of G_N_ and G_C_, two forms of TSWV G_N_, from nucleotide 109-1353 (named G_N_-1353) and 109-1386 (G_N_-1386), as well as both soluble forms, G_N_S (109-930) and G_C_S (1354-3174) were cloned into pBT3-SUC and pPR3N and used for interaction test. The MbY2H analysis revealed self-interactions of the TSWV NSs, N, NSm, G_N_S, G_N_-1353 and G_C_ regardless of the MbY2H vector used (Figure 4). A G_N_S self-interaction indicated that the ability of G_N_ to dimerize resides in the predicted ectodomain. Homodimerization of N and NSm was also observed using various expression systems (Uhrig et al., 1999; Leastro et al., 2015). The self-interaction of NSs was only observed in another tospovirus, CaCV (Widana Gamage and Dietzgen, 2017). We also detected interactions among different forms of G_N_ and G_C_, including G_N_S-Cub and NubG-G_N_-1386, G_N_-1353-Cub and NubG-G_N_-1386 as well as G_C_-Cub and NubG-G_N_-1353. In addition to the self-interaction, G_C_-Cub also interacted with NubG-G_N_-1353, NubG-N and NubG-NSs (Figure 4).

**Figure 4.**
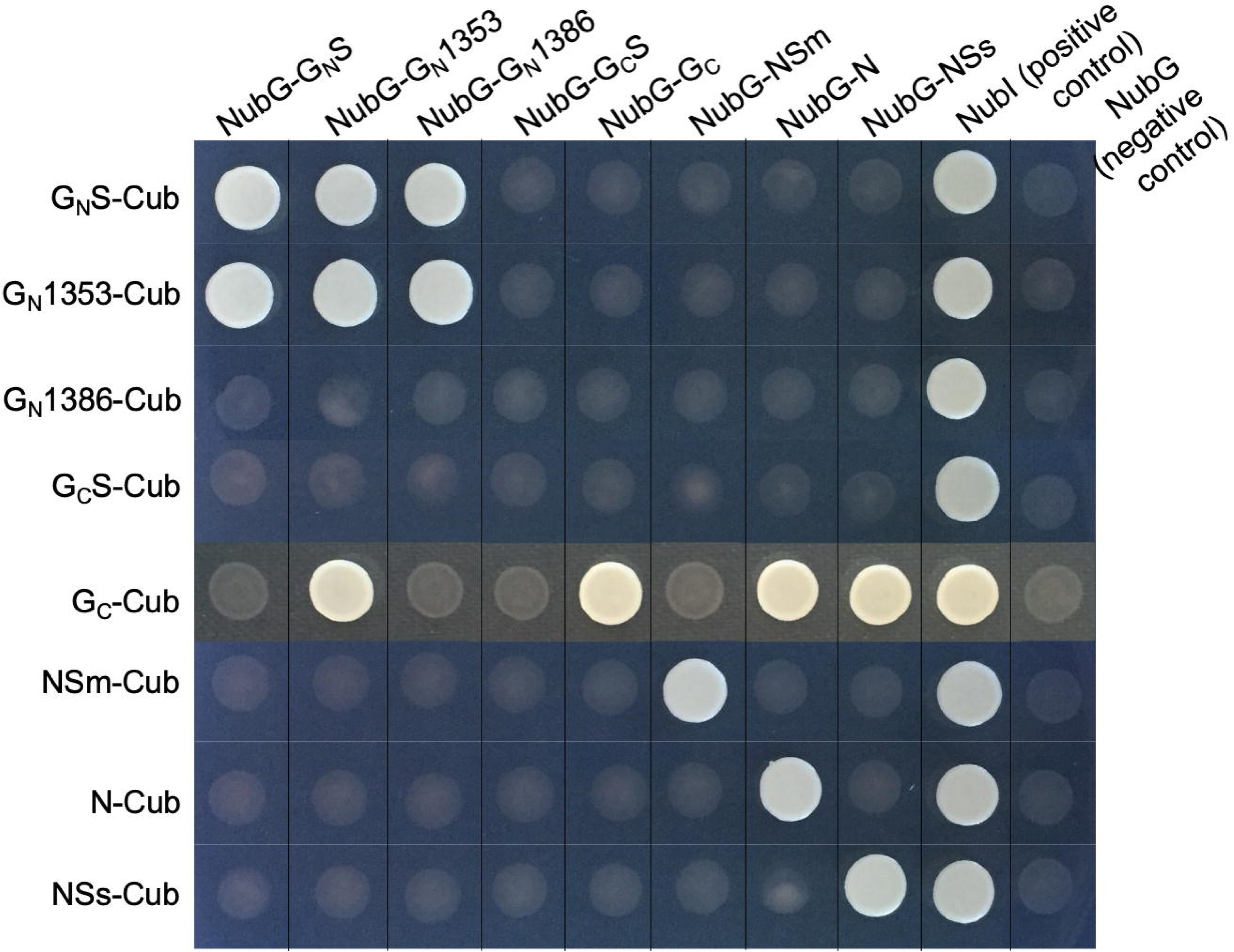
Validation of interactions between TSWV proteins using a split-ubiquitin membrane-based yeast two hybrid (MbY2H) assay. Each TSWV protein was expressed as TSWV-Cub (pBT3-SUC) and NubG-TSWV (pPR3N). Recombinant plasmids pBT3-SUC-TSWV were transformed into NYM51 yeast competent cells first to express TSWV-Cub, then each pPR3N-TSWV plasmid was transformed to NYM51 cells expressing individual TSWV-Cub. Interactions between TSWV protein interactions were determined by the interactions between fusion proteins TSWV-Cub and NubG-TSWV. The transformants were grown on QDO media, and interactions of GC-Cub and NubG-TSWV were validated on the QDO media supplemented with 100mM 3-AT. Interactions between TSWV-Cub and NubI, and between TSWV-Cub and NubG were used as positive and negative controls. Co-transformation of pTSU2-APP and pNubG-Fe65 into NYM51 was used as another positive control (data not shown). QDO, yeast quadruple-dropout (SD/–Ade/–His/–Leu/–Trp) medium.

## Discussion

Tomato spotted wilt virus was first characterized in 1919 (Brittlebank, 1919), and is the best characterized virus within the orthotospovirus genus., However, there remain many unanswered questions regarding how this virus functions as it infects both plant and animal cells. The structure of the virus is conserved in both cell types, with a genome coated with a capsid protein (N) which then buds through a host membrane embedded with viral glycoproteins G_N_ and G_C_ (for review see (Oliver and Whitfield, 2016)). Due to this conservation of the structure, the hypothesis guiding this study was that the localization patterns of the proteins would be conserved between plant and insect cells. This was supported by a previous study that G_N_ and G_C_ had localization patterns that were conserved in BHK21 cells (not a host to this virus)(Kikkert et al., 2001). Previously another virus which infects also both plants and insects (Maize mosaic virus, MMV, alphanucleorhabdovirus), also demonstrated that protein localization is conserved in the divergent hosts (Martin and Whitfield, 2018).

Like MMV, although there was a significant amount of conservation between the two cell types, NSm has a divergent localization pattern when comparing cells. In both systems, in plants, the movement proteins demonstrate a localization pattern consistent with the plasmodesmata of the cell (Martin and Whitfield, 2018, this study). This pattern in plants was expected for NSm as it matches previous studies (Kormelink et al., 1994; Li et al., 2009). For insects, we expected to see something that varied from MMV because NSm, although also in 30K superfamily of plant movement proteins similar to MMV-3, forms tubules to assist in movement, and 3 does not (Melcher, 2000; Li et al., 2009). This prediction proved accurate, with NSm in insect cells forming larger aggregations compared to MMV-3 (Martin and Whitfield, 2018). The localization in the cytoplasm itself was not surprising as the possibility exists that NSm may function to move either viral proteins or viral complexes around the cell during the course of infection similar to the rhabdoviruses (Min et al., 2010). This is further supported by the interaction between NSm and Dnaj which may assist in intracellular movement through an HSP-70 dependent mechanism (Soellick et al., 2000). The localization pattern between MMV-G and TSWV-G_N_ and G_C_ also showed differences as well that were expected, as MMV is a alphanucleorhabdovirus, it shows a strong localization pattern in the membranes surrounding the nucleus, even in insect cells (Martin and Whitfield, 2018). In TSWV, the TSWV-G_N_ and G_C_ localization patterns are consistent in insects with an ER and Golgi association that is also seen in the plant cells, both intact cells and protoplasts (Ribeiro et al., 2008).

Based on the conservation between insect and plant cells, it was then of interest to determine the localization pattern in infected cells in comparison to healthy cells. At the onset of the study, we expected that the endoplasmic reticulum would change during infection based on previous work, giving us a marker to determine infection status (Ribeiro et al., 2008). However, a RFP-ER transgenic line (Martin et al., 2009) showed no obvious changes in the cells during infection. Other possible markers were tried, however, unfused RFP, which has a cytoplasmic and nuclear localization, was best. In these plants, darker areas were observed where the RFP was excluded which did not occur in healthy plants indicating a possible site of viral protein accumulation. This was reminiscent of earlier work with the nuclear marker and Sonchus yellow net virus, a betanucleorhabdovirus, which also shows a similar exclusion at areas where virions/viral proteins accumulate (Goodin et al., 2007). With an established marker, we expected to see differences in the localization patterns during infection as previous data for G_C_ indicated that this protein changes localization when G_N_ and N are present (Ribeiro et al., 2009).

As seen in Figure 2, we observed a dramatic difference in G_C_ when expressed in the presence of viral infection matching previous data (Ribeiro et al., 2009). However, the visualization of N, NSs and NSm surrounding the voids present with RFP was interesting. It was expected that N would be present as it would interact with the viral RNA during morphogenesis and previous data had indicated that N moved in the cell over time from an outer to inner localization pattern (Montero-Astúa, 2012) NSm may function in this case to move N and other viral proteins from the cell periphery where it is originally observed to the site of maturation. This is also consistent with the rise in the expression levels of NSm observed previously (Kormelink et al., 1994) and our own observation of a movement of NSm from the cell periphery to the interior of the cell over time (Supplementary figure 1). It is also consistent with NSm having weak RNA binding capacities which may assist in cellular movement (Soellick et al., 2000). The role of NSs and its movement over time similar to N and NSm (Supplemental figure 2) suggests that the role of NSs as a silencing suppressor (Takeda et al., 2002) may require the movement to the areas where the viral genome accumulates or that NSs may have other roles during infection that have not been identified.

To further support the localization patterns consistent with the viral structure, the interactions between viral proteins were explored. Initially, we expected to see an interaction between G_N_, G_C_ and N (Snippe et al., 2007; Ribeiro et al., 2009), an interaction between N and NSm (Soellick et al., 2000), and an N-N interaction (Uhrig et al., 1999; Snippe et al., 2005). We selected bimolecular fluorescence complementation (BiFC) to test for interactions as it had been successfully used in three other orthospoviruses (Dietzgen et al., 2012; Tripathi et al., 2015; Widana Gamage and Dietzgen, 2017). We confirmed an N-N and and NSm-NSm interaction in TSWV that is also seen in all other tested orthotospoviruses and suggests these may be conserved across the genus (Figure 5) (Dietzgen et al., 2012; Tripathi et al., 2015; Widana Gamage and Dietzgen, 2017). We also observed new interactions, such as NSs and Gc and N and Gc which may be part of the virion assembly process. These interactions demonstrated a temporal effect as well, being more detectable over background at particular time points consistent with the changing localization patterns for N, NSs and NSm observed previously. However, we did not see an N-NSm interaction that was previously reported using protein expression and overlay assays (Soellick et al., 2000). This was confusing as this interaction was demonstrated in Impatiens necrotic spot virus (INSV) and Iris yellow spot virus (IYSV) using BiFC (Dietzgen et al., 2012; Tripathi et al., 2015) and is considered important to ferry a ribonucleoprotein complex from cell-to-cell through the plasmodesmata (Kormelink et al., 1994; Soellick et al., 2000; Leastro et al., 2017). Perhaps in this case, the interaction may occur when using other techniques and the possiblity that the YFP halves in our case may block the possible interaction. This was further explored using yeast two-hybrid (Y2H).

**Figure 5.**
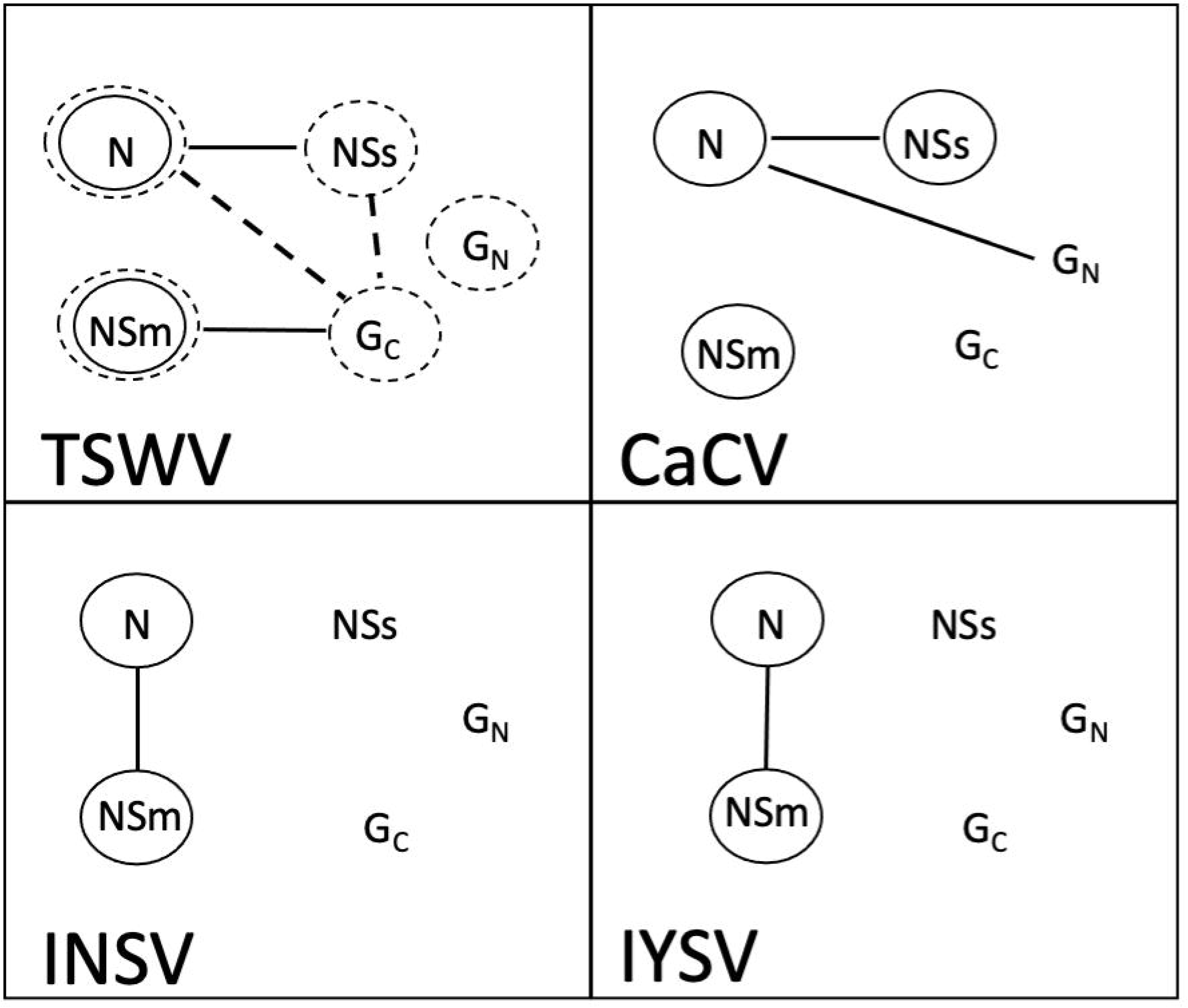
Interaction maps of characterized orthotospoviruses. The top left panel is the interaction map of tomato spotted wilt virus. The second panel on the top are the interactions shown for Capsicum chlorosis virus (Widana Gamage and Dietzgen, 2017). The bottom left panel are the interactions seen in Impatiens necrotic spot virus. (Dietzgen et al., 2012) The bottom right panel are the interactions seen in Iris yellow spot virus. (Tripathi et al., 2015) Circles indicate protein self-interactions and lines between proteins are non-self-interactions. Solid lines mark interactions that were demonstrated through bimolecular complementation and those that are dashed were demonstrated through yeast-two hybrid analysis.

In Y2H, we were able to confirm the N-N and NSm-NSm interaction but did not detect the N-NSm interaction. As two different methods were used, the conclusion became that perhaps viral strain differences or the tags added to the proteins cause the discrepancy. Previously, TSWV strain L3 was utilized to test for N-NSm (Soellick et al., 2000) and our studies were conducted exclusively with TSWV MT2. Looking at an alignment between L3 and MT2 N and NSm proteins, there are changes in the amino acids, with four in N and ten in NSm. There are a few possibilities to consider: perhaps TSWV may have different interactions depending on the strain in question. With a global distribution and a large host range, it may suggest that adaptations to hosts results in changes in the interaction maps depending on strains. It is also possible that the interaction in MT2 is facilitated by a host factor and is a direct interaction in L3. The other possibility concerns experimental design, we did not conduct in vitro interaction assays with SDS gels and detection of interactions through antibody binding, but instead relied on the use of tags both in Y2H and BiFC and this may suggest that the presence of a larger tag may block the interaction between these proteins in TSWV which did not occur with a his tag used previously (Soellick et al., 2000). We also did not detect the N-Gc or the NSs-Gc interaction in yeast which could potentially be accounted for by the changes over time that can be observed in plants which are not seen in insect cells suggesting this is a plant specific response.

The discovered interactions of TSWV and other orthotospoviruses proteins are important for deciphering the biological roles these viral proteins play in the cells of its host plants and insects. The next step in fully understanding the TSWV-thrips-plant host system is the identification of the host proteins that assist in viral replication, movement, or other steps in the infection cycle. These host proteins represent new targets for controlling orthotospoviruses in either insects (Maurastoni et al., 2023) or plants (Hsu and Spindler, 2012; Akhter et al., 2021). Identification of host protein involvement may also help to resolve the detected (and non-detected) interactions of TSWV-MT2 to a larger degree and to determine the roles of these interactions during infection. Resistance can also be achieved by manipulating these host factors in either host. This would be an environmentally friendly strategy to achieve virus control that would in turn increase agricultural productivity and reduce the use of toxic chemicals aimed at controlling thrips vectors.

---

Supplemental Figure 1: Localization of TSWV proteins over time. Left side) TSWV proteins expressed in plant cells at 2 days post infection. Column 1, GFP tagged to TSWV protein, Column 2, RFP, Column 3, overlay of columns 1 and 2. Rows are as follows: I-III, TSWV-N-GFP, IV-VI, TSWV-NSs-GFP, VII-IX, TSWV-GFP-NSm. *Right side)* TSWV proteins expressed in plant cells at 3 days post infection. Column 1, GFP tagged to TSWV protein, Column 2, unfused RFP, Column 3, overlay of columns 1 and 2. Rows are as follows: X-XII, TSWV-N-GFP, XIII-XV, TSWV-NSs-GFP, XVI-XVIII, TSWV-GFP-NSm.

Supplemental Figure 2: Initial transformation of yeast vectors plated on double dropout media (DDO SD/–Leu/–Trp). Each TSWV protein was expressed as TSWV-Cub (pBT3-SUC) and NubG-TSWV (pPR3N). Recombinant plasmids pBT3-SUC-TSWV were transformed into NYM51 yeast competent cells first to express TSWV-Cub, then each pPR3N-TSWV plasmid was transformed to NYM51 cells expressing individual TSWV-Cub. Double-dropout media was utilized to determine that each combination could express well to validate the quadruple dropout media results.

Supplemental Video 1: Localization of TSWV-N-GFP in infected plant cells using Z stacks to scan through cell layers. GFP is tagged to the nucleocapsid protein, while RFP is expressed alone. The white scale bar indicates 10µm.

Supplemental Video 2: Localization of TSWV-NSs-GFP in infected plant cells using Z stacks to scan through cell layers. GFP is tagged to the non-structural silencing protein, while RFP is not fused to any protein. The white scale bar indicates 10µm.

Supplemental Video 3: Localization of TSWV-NSm-GFP in infected plant cells using Z stacks to scan through cell layers. GFP is tagged to the non-structural movement protein, RFP is expressed as a non-fused protein. The white scale bar indicates 20µm

Supplemental Video 4: Localization of TSWV-G_C_-GFP in infected plant cells using Z stacks to scan through cell layers. GFP is tagged to the glycoprotein G_C_, while RFP is expressed as a non-fused protein. The white scale bar indicates 10µm

