## Supplemental Figure 2 for "Visualizing tomato spotted wilt virus protein localization: Cross-kingdom comparisons of protein-protein interactions"

### Slide 1
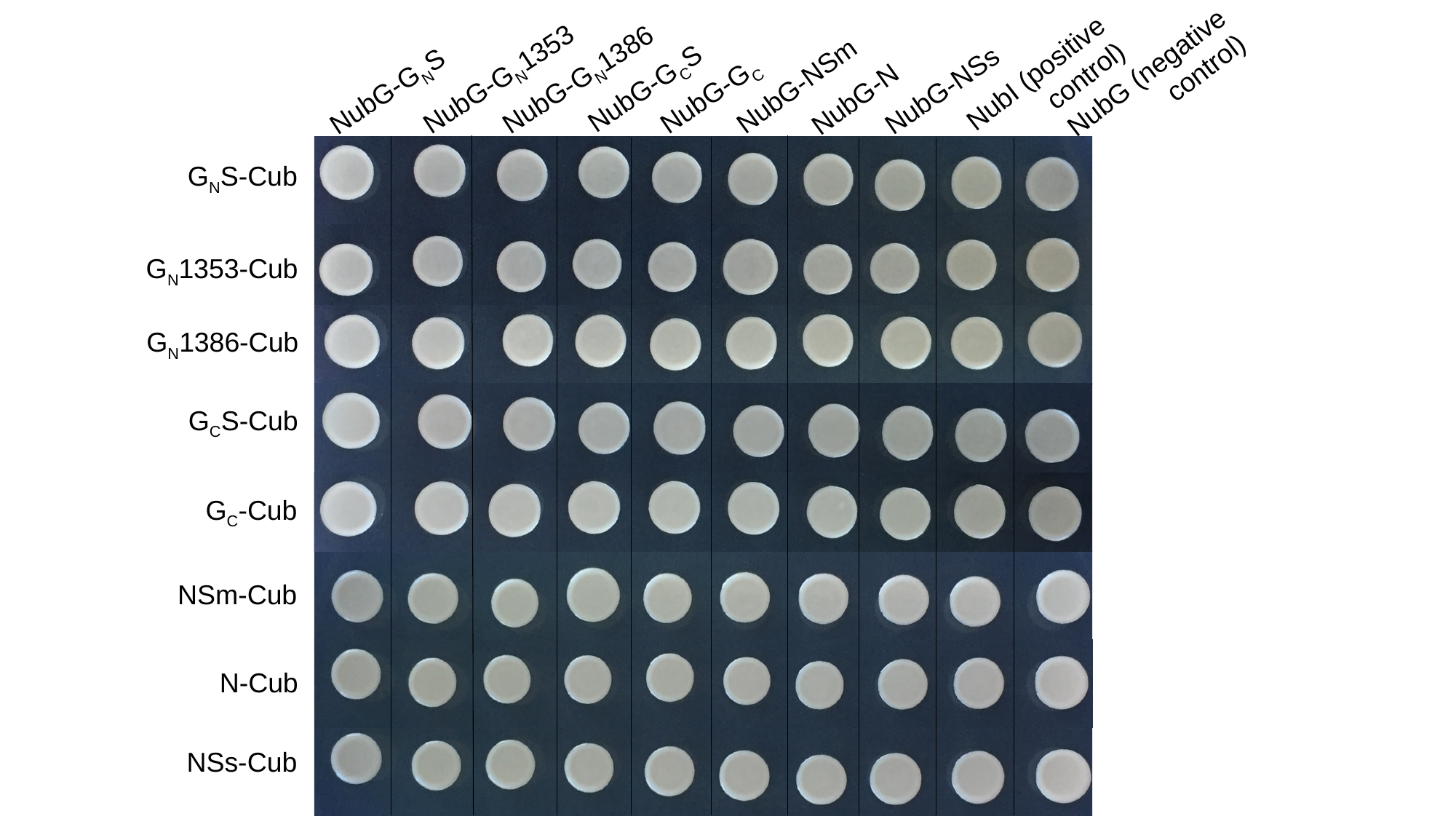

NubG-G_N_1353
NubG-G_N_1386
NubG-G_N_S
NubG (negative control)
NubI (positive control)
NubG-N
NubG-NSm
NubG-NSs
NubG-G_C_S
NubG-G_C_
G_N_S-Cub
G_N_1353-Cub
G_N_1386-Cub
G_C_S-Cub
G_C_-Cub
NSm-Cub
N-Cub
NSs-Cub
