## Supplementary figures and images for "Visualizing tomato spotted wilt virus protein localization: Cross-kingdom comparisons of protein-protein interactions"

### Supplemental Figure 1

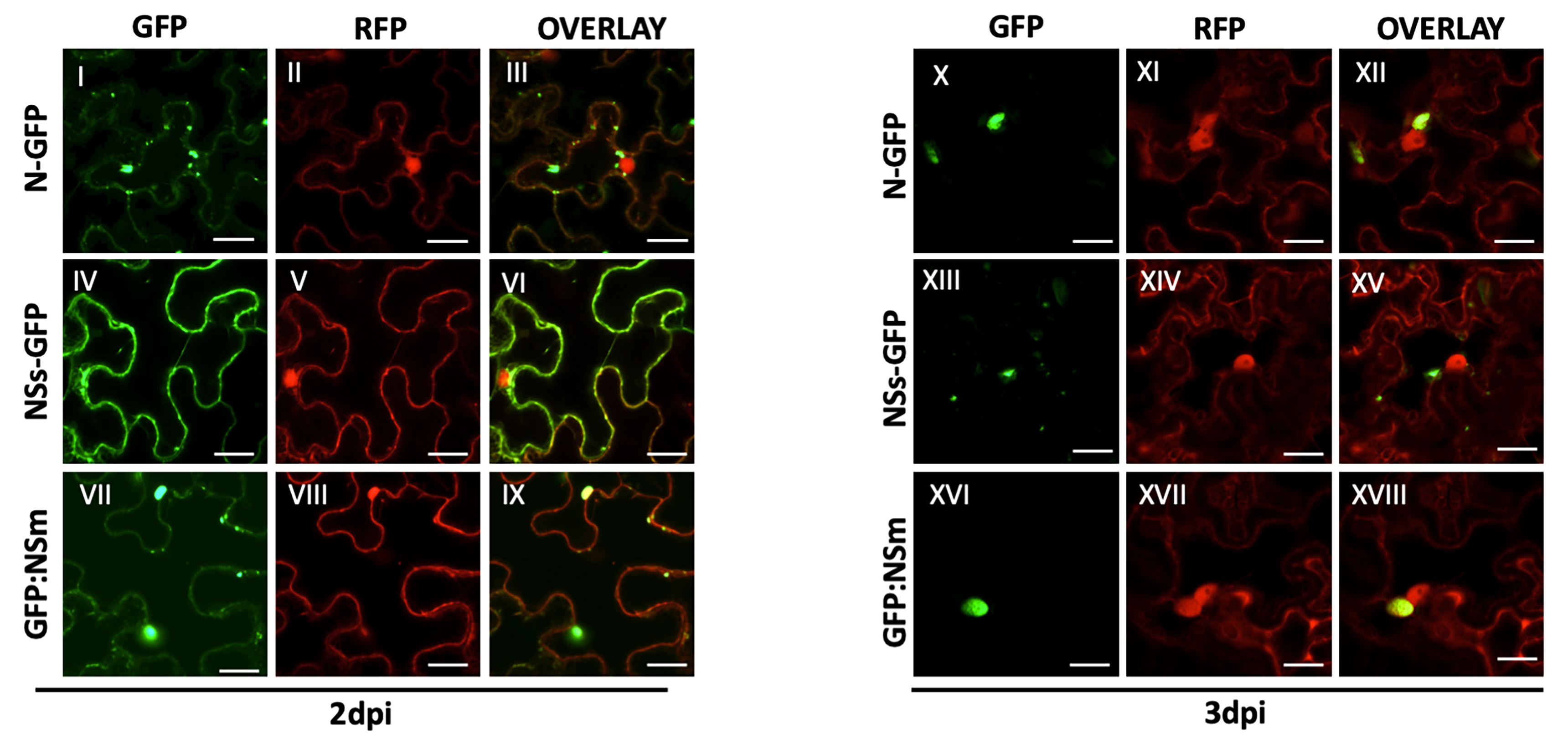
